# An Open-Access Database of Video Stimuli for Action Observation Research in Neuroimaging Settings: Psychometric Evaluation and Motion Characterization

**DOI:** 10.1101/2023.11.17.567513

**Authors:** Christian Georgiev, Thomas Legrand, Scott J. Mongold, Manoa Fiedler-Valenta, Frédéric Guittard, Mathieu Bourguignon

## Abstract

Video presentation has become ubiquitous in paradigms investigating the neural and behavioral responses to observed actions. In spite of the great interest in uncovering the processing of observed bodily movements and actions in neuroscience and cognitive science, at present, no standardized set of video stimuli for action observation research in neuroimaging settings exists. To facilitate future action observation research, we developed an open-access database of 135 high-definition videos of a male actor performing object-oriented actions. Actions from 3 categories: kinematically natural and goal-intact (*Normal*), kinematically unnatural and goal-intact (*How*), or kinematically natural and goal-violating (*What*), directed towards 15 different objects were filmed from 3 angles. Psychometric evaluation of the database revealed high video recognition accuracy (*Mean* accuracy = 88.61 %) and substantial inter-rater agreement (Fleiss’ *Kappa* = 0.702), establishing excellent validity and reliability. Videos’ exact timing of motion onset was identified using a custom motion detection frame-differencing procedure. Based on its outcome, the videos were edited to assure that motion begins at the second frame of each video. The videos’ timing of category recognition was also identified using a novel behavioral up-down staircase procedure. The identified timings can be incorporated in future experimental designs to counteract jittered stimulus onsets, thus vastly improving the sensitivity of neuroimaging experiments. All videos, their psychometric evaluations, and the timing of their frame of category recognition, as well as our custom programs for performing these evaluations on our, or on other similar video databases, are available at the Open Science Framework (https://osf.io/zexc4/).

## Introduction

In the past two decades, action observation research has been facilitated by technological advancements which allow researchers to easily record and present videos of biological motion. This in turn allowed for the use of video stimuli in controlled neuroimaging and psychophysics investigations of the neural processing of the movements which make up human naturalistic action. Consequently, experimental paradigms involving presentation of pre-recorded videos of various types of bodily motion have become ubiquitous in neuroscience, cognitive science, and psychology. Such investigations have produced great advances in the understanding of biobehavioral phenomena ranging from the cellular to the cognitive level, such as the activity of mirror neurons (Arnstein *et al*., 2011; Braadbaart, Williams and Waiter, 2013; Moriguchi *et al*., 2009), the Mu and Beta oscillations of the sensorimotor cortex (Avanzini *et al*., 2012; Brunsdon, Bradford and Ferguson, 2019; Muthukumaraswamy and Johnson, 2004; Quandt and Marshall, 2014), motor affordances (Bach, Bayliss and Tipper, 2011; Tipper, Paul and Hayes, 2006), action identification and discrimination (Orban, Ferri and Platonov, 2019; Urgen and Orban, 2021; Vannuscorps and Caramazza, 2016), gesture recognition (Beattie and Shovelton, 2002), observation learning (Buccino *et al*., 2004; Malfait *et al*., 2010), theory of mind (Caillaud *et al*., 2020; Saxe *et al*., 2004; Sylwester *et al*., 2012), and empathy (Pütten *et al*., 2014; Tholen *et al*., 2020). To address these phenomena, typically, videos of an agent performing a manual movement are presented to observers either in blocked or in pseudo-random counterbalanced designs while electrophysiological or hemodynamic measures of neural activity are recorded. This approach allows for a resolute assessment of the neurocognitive processing of observed actions, carried out by bottom-up perceptual as well as top-down cognitive neuronal mechanisms. Thus, the approach allows for the investigation of the underpinnings of perception, comprehension, and acquisition of complex action and intention in both neurotypical (Biagi et al., 2016) and neurodivergent (Scott *et al*., 2020; Spengler, Bird and Brass, 2010) populations.

To effectively capture the neural correlates of the detection and comprehension of the nuanced movements which comprise complex naturalistic action, however, the presented video stimuli must be constructed in accordance with the highest psychometric standards. Duration, frame rate, pixel resolution, color palette, luminosity, filming angle, depicted body segments, gender and handedness of the actor, familiarity with the action and the actor, and recognizability of the action have all been controlled for, at least to a certain extent, in many individual action observation studies. Yet, across studies, there is substantial heterogeneity in the presented video stimuli, with some authors presenting videos depicting only the hand, others presenting videos depicting the entire upper body of the actor; some authors presenting videos from a first-person perspective, others from a third-person perspective; and some authors presenting videos in black and white, others in color. With the advancement of video recording equipment, the overall quality of the video stimuli increases and novel tools have been developed for monitoring changes across video recordings, including nuanced changes in posture (Zouba *et al*., 2008) or motion kinematics (Trettenbrein and Zaccarella, 2021) of the actor across videos. Still, standardized sets of high-quality stimuli, made specifically for action observation research in neuroimaging settings, which could be used across experiments and laboratories, are scarce, and most investigators in the field construct their own stimuli or rely on non-dedicated freely-available videos from the internet. The reliance on such stimuli which likely differ in either technical properties (e.g., pixel resolution and frame rate) or higher-order characteristics of motion (e.g., goal-directedness and intention relation) prevents the comparison of results across experiments, complicates the conduction of robust and reliable meta-analyses, and hence negatively impacts the reproducibility of action observation research. More recently, open-access multipurpose databases of video stimuli have been developed (Di Crosta *et al*., 2020; Umla-Runge *et al*., 2012; Urgen *et al*., 2022), however, videos therein feature only kinematically correct natural actions and were not characterized in terms of their action onset which complicates their use in neuroimaging settings.

An initial challenge of the serial presentation of a large number of videos in neuroimaging settings is that differences in the timing of movement onset can exist across videos, even among the videos of a carefully constructed stimulus set where all videos are controlled for duration and frame rate. Although some authors have identified and reported the timing of motion onset of their stimuli (e.g., Platonov and Orban, 2016), this practice is often overlooked or underreported in the literature. This is especially an issue for electrophysiological neuroimaging experiments since these typically require averaging of neural responses evoked by a large number of videos (Huettel, 2012), and both time domain and time-frequency domain analyses have to be precisely timed to video onset. In that setting, seemingly small differences in the actual visual detection of movement across videos are likely to contaminate the contrasts between the experimental conditions. For instance, if an onset trigger is sent at the very first frame of each video stimulus’ presentation, a discrepancy in movement onset across videos of as little as 3 frames at 60 Hz translates to a jitter of 50 ms. Such jitter can substantially impact the shape and timing of the observed evoked neural responses, resulting in noisier and flattened out averaged neural responses and hence, difficulties in detecting differences between experimental conditions. In hemodynamic neuroimaging, the negative impact of discrepancies between video onset and actual movement onset also exists, although it may be smaller due to the slow nature of the blood-oxygen-level-dependent (BOLD) signal.

In an analogous fashion, the issue of timing and triggering also concerns the timing of the actual conscious recognition of the higher-order properties of movement, such as the degree of integrity of movement kinematics, goal, or intention across different videos, which are the central manipulation of numerous studies addressing the neural and cognitive representations of normal and abnormal movement (e.g., Stapel *et al*., 2010; Cheng *et al*., 2017). Such studies often imply presentation of videos that either depict natural or unnatural actions with varying degrees of goal integrity. The videos of natural actions typically comprise kinematically correct movements, as would normally be performed and observed in everyday settings, whereas the videos of unnatural actions comprise kinematically incorrect odd movements, which are unlikely to be performed or observed in everyday settings. However, the motor redundancy and abundance of any complex human action (Gera *et al*., 2010; Steen, van der Steen and Bongers, 2011) suggest that it is highly likely that the kinematic nature and goal integrity of the actions in different videos are recognized at very different time points. Once again, averaging neural responses with jittered onsets will impact the shape and timing of the observed evoked neural responses and hence, severely contaminate any contrast between natural and unnatural, and between goal-intact and goal-violating action categories.

Therefore, determining when within a video stimulus movement onset and kinematic and goal category are actually perceived is crucial for effective experimental design. Although precise motion characterization of videos can be obtained with complex artificial intelligence computer vision algorithms (Vrigkas *et al*., 2015), such algorithms are not always freely available and require advanced expertise in deep learning. With respect to motion onset detection, more user-friendly motion detection systems based on the frame-differencing method have been widely applied on video recordings within the field of counseling psychology (Paxton and Dale, 2013; Ramseyer and Tschacher, 2011), however, no such methodology has yet been applied for characterizing videos used as stimuli in action observation research. We propose that such a system can successfully be applied for identifying the first frame of motion onset within video stimuli, so that all video stimuli can subsequently be edited to start exactly at the identified frames. With respect to the recognition of higher-level properties of categories of videos with different kinematics and goal-integrity, no accessible and user-friendly systematic stimulus evaluation procedure exists. We propose that a straightforward psychophysics approach based on the classic up-down staircase methods (Levitt, 1971; Kaernbach, 1991; Wetherill and Levitt, 1965) can be applied for identifying the first frame at which a video can be categorized (i.e., recognition of the movement as kinematically correct or incorrect and goal-intact or goal-violating). Namely, if a video is played up until a given number of frames, then paused, and an observer is asked whether the motion depicted in the video was kinematically correct or not, and goal-intact or not, their response can be used for adjusting the number of frames played on the subsequent presentation of the same video to another observer. With enough between-subject iterations, an asymmetric staircase procedure governed by two simple rules: 1) If the category of motion has been correctly detected within the played segment, the number of played frames decreases on the subsequent presentation and 2) If the category of motion has not been detected within the played segment, the number of played frames increases on the subsequent presentation; should converge on the frame where the category of the video becomes discernible for naïve human observers. The identified frames can be used for implementing precise triggering for the identification of brain or behavioral responses to action kinematics and goal integrity.

Aspiring to improve the feasibility of action observation research, we introduce a large psychometrically evaluated open-access database of videos tailored to the demands of neuroimaging experiments. More precisely, we present 135 videos depicting a goal-directed action that is either kinematically correct or not, and goal-intact or not, filmed from a third-person perspective from 3 angles (portrait, left profile, and top). These videos should be useful for future investigations of the neural mechanisms of action and goal perception and comprehension. Moreover, we provide an easy to implement, open-source frame-differencing motion detection system for identifying the frame of motion onset within each video and a straightforward procedure for editing the videos based on their motion onset. We also provide a straightforward open-source up-down staircase procedure for identifying the frame on which kinematics and goal-integrity are recognized within each video. Both procedures are purposefully designed in a way that would allow any reader to easily implement them on our, or another set of similar video stimuli that depict motion of a single agent.

## Methods

### Participants

Fifty-one adults (27 Female, *Mean* ± *SD* age: 28.47 ± 7.16 years) provided ratings for psychometric evaluation of the videos. A different sample of 17 adults (8 Female, *Mean* ± *SD* age: 25.82 ± 4.6 years) participated in the up-down staircase category recognition procedure. Participants were recruited from the Université Libre de Bruxelles (Brussels, Belgium). All participants were healthy, with no known neurological or psychiatric disorders, had normal or corrected-to-normal vision, and gave their written consent for participation in the study. The protocols were approved by the local ethics committee and the study was conducted in accordance with the Declaration of Helsinki.

### Stimuli

A database of videos was created specifically for the demands of subsequent action observation experiments. The database consists of high-resolution video recordings filmed in 4K (at 60 frames/second with a 12-megapixel Apple iPhone 12 camera in portrait orientation, positioned on a stationary tripod) and subsequently resized to 720 × 1280 pixels. The videos depict a male actor (Caucasian, Age = 22 years) seated in front of a table and performing a goal-directed action with a common everyday object (calculator; cap; coffee jar; comb; computer mouse; cup; fork; glasses; hat; headphones; hourglass; pen; pencil case; ruler; scissors). With respect to kinematic and goal integrity, actions from 3 categories were filmed: *Normal* (i.e., the object was used in the most kinematically natural way and for its intended goal; Figure 1A), *How* (i.e., the object was used in a kinematically unnatural way and for its intended goal; Figure 1B), and *What* (i.e., the object was used in a kinematically natural way but not for its intended goal; Figure 1C). Inspired by Iacoboni *et al*. (2005) and following the recommendations of Sonkusare, Breakspear and Guo (2019), all actions were filmed in living settings, providing a static naturalistic background and hence increasing the ecological validity of the stimuli. Each action was filmed from a third-person point of view from 3 angles: portrait, left profile, and top (Figure 2). In total, 15 actions were filmed in each category and from each filming angle resulting in 135 videos, each with a duration of 3–6 s. All videos in .m4v and .mat format can be found in folder “Video Database” at the Open Science Framework: https://osf.io/zexc4.

**Figure 1.**
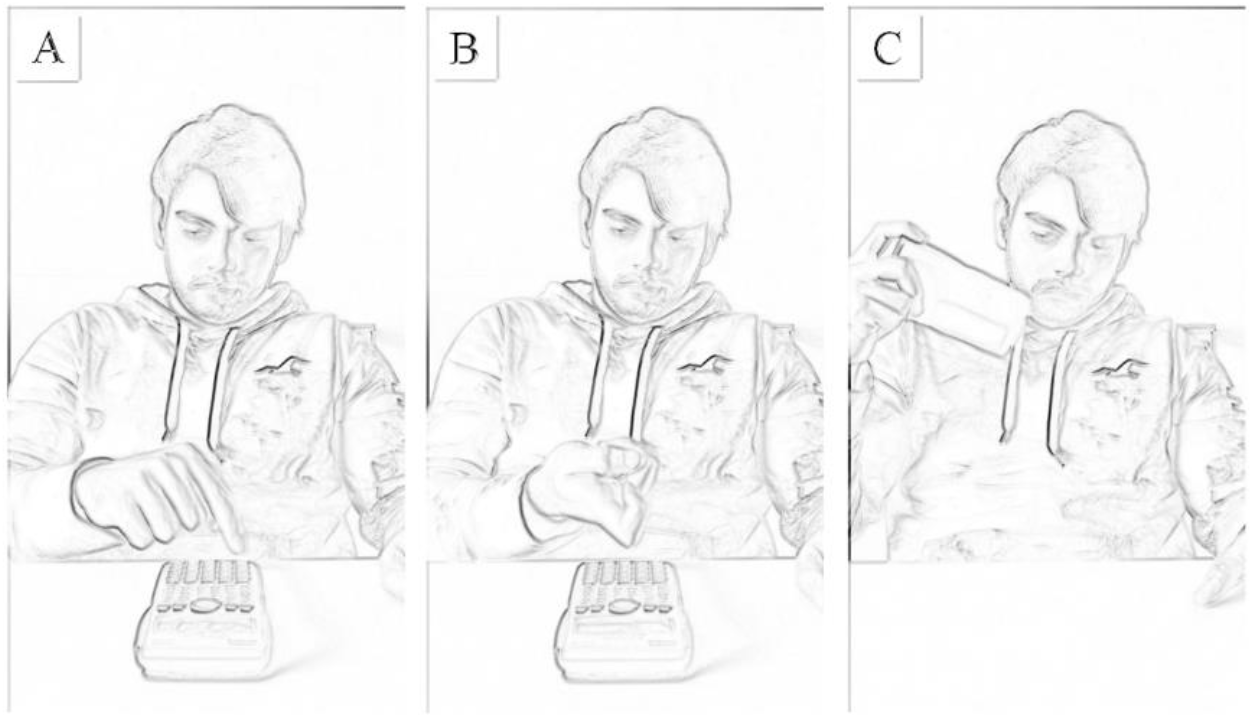
^1^ Representative frames extracted from a video depicting a Normal (A), How (B), and What (C) use of a calculator.

**Figure 2.**
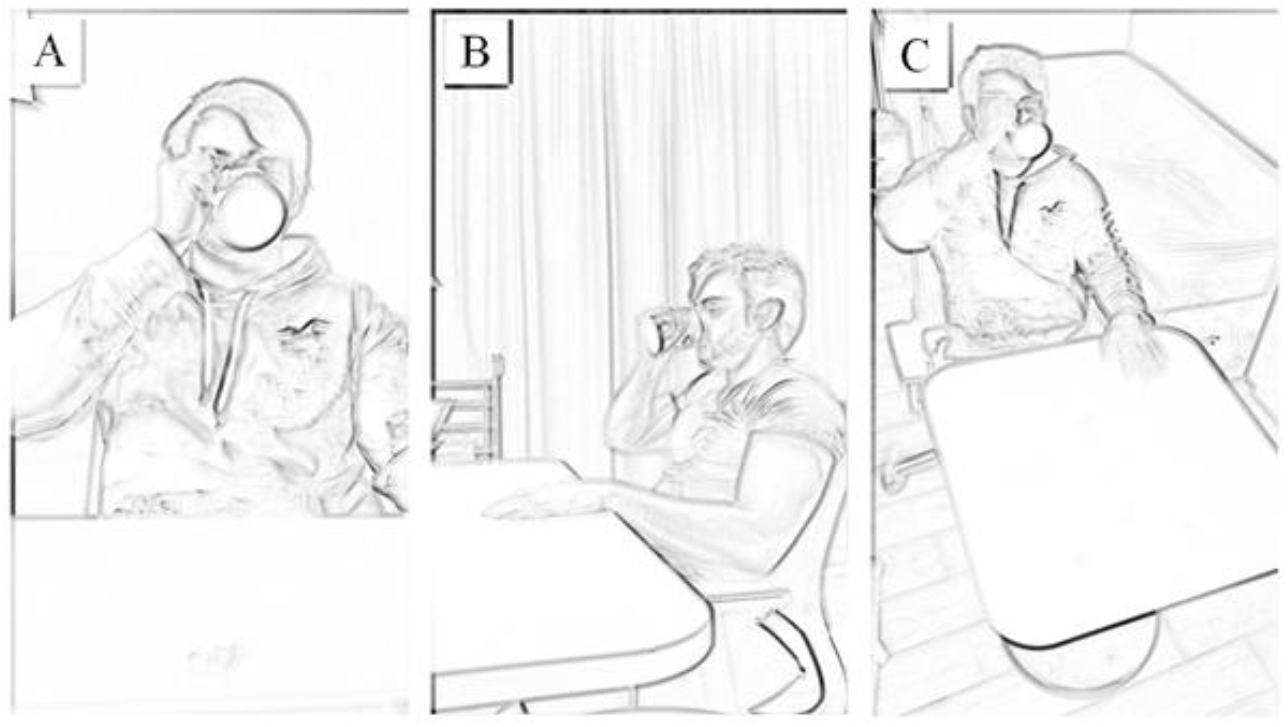
^1^ Representative frames extracted from videos depicting a Normal use of a cup from a portrait (A), profile (B), and a top angle (C).

### Psychometric Evaluation

The participants were instructed that they would be asked to evaluate a number of videos in terms of their kinematic and goal integrity category. The 3 categories of videos, along with their labels (*Normal, How*, or *What*) were explained to the participants and they were allowed to ask for further clarification if necessary. The participants were then seated in front of a 17-inch computer monitor and were sequentially presented with the 135 videos in a pseudo-random order (multiple presentations of the same video were not allowed). After the end of each video, participants were prompted to classify it as “*Normal*”, “*How*”, or “*What*” by pressing a corresponding key on the keyboard. The entire rating procedure was completed in a single session, with no time constraint.

### Identification of the Frame of Motion Onset

All videos were passed as input to a sensitive custom-made motion detection system based on the OpenCV library (Bradski and Kaehler, 2008) in Python 3 (Python Software Foundation, Wilmington, DE). This system was made such that it detected and reported the presence of motion per each frame of the input video object (Figure 3). To this end, each frame of a video was captured, converted to grayscale, and compared to the static initial frame of the respective video using a frame-differencing algorithm (Ramseyer, 2020).

**Figure 3.**
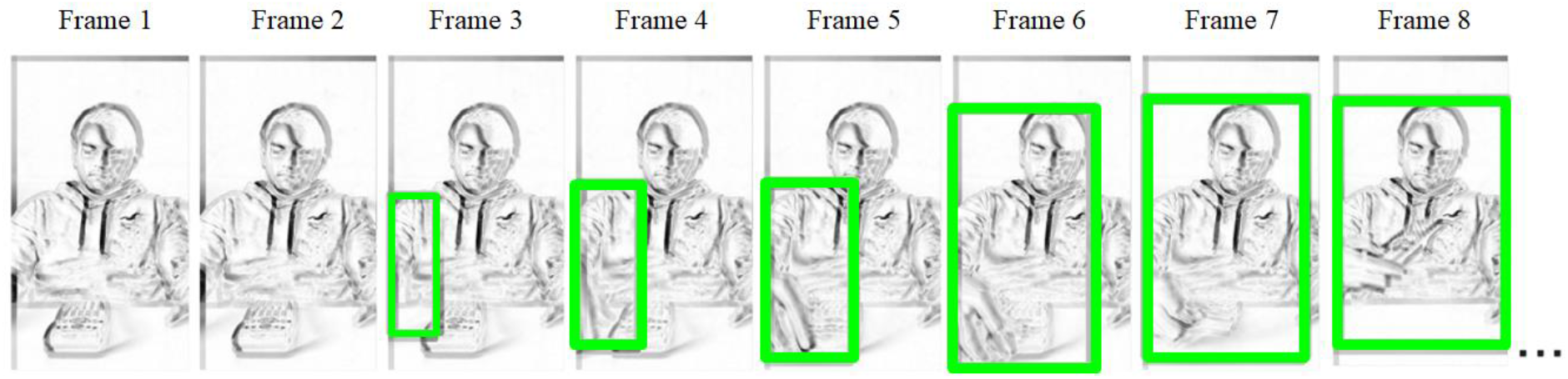
^1^ Schematic illustration of the operation of the motion detection system.

Differences between the two frames were highlighted with a binary threshold method, setting pixels that were at least 10 % different on the grayscale to black and the rest to white. Subsequently, the contours of the resulting binary frame were extracted using a standard border following algorithm (Suzuki and Abe, 1985) and used to perform box detection of motion. To avoid detecting background noise as motion, a threshold on the minimal box area was set to 400 pixels and restricted in the Y dimension to the area corresponding to the location of the actor, thus assuring the detection of only meaningful biological motion of the actor. The first frame on which the system registered a difference between the two frames is reported as the frame of motion onset. For each video, the outcome of the frame-differencing procedure was inspected and compared to the frame of motion onset identified by a human observer (C. G.) during a frame-by-frame video presentation to assure accurate identification and reporting of each video’s frame of motion onset. In case of disagreement, a correction was applied in favor of the judgment of the human observer. Video stimuli were then edited to start at the frame preceding that of motion onset. The Python scripts for performing the frame-differencing motion onset detection on this or any other similar set of videos and the script for breaking the videos into frames can be found in the folder “Motion Onset Frame-Tracing Procedure” at https://osf.io/zexc4/.

### Identification of the Frame of Category Recognition

Following-up the general video recognition procedure, we implemented a novel approach for identifying the frame on which videos became discernibly recognizable as *Normal, How*, or *What* to human observers. The experimental settings and the instructions given to the participants were identical to the ones for the psychometric evaluation procedure. Each participant completed a single iteration of an up-down staircase implemented by a custom-made MATLAB (Mathworks, Natick, MA) script. For each iteration, all videos were played once in a pseudo-random order, up to a specific frame. For the first iteration of the staircase, performed by the first participant, each video was played up to the frame corresponding to the 50th percentile of the video’s frames and then paused. After each video’s pause, the participant was prompted to report whether, based solely on the played video segment, they would classify the given video as “*Normal*”, “*How*”, or “*What*” via mouse click. On each subsequent iteration, performed by each subsequent participant, the number of frames that would be played from each video was adjusted based on the response of the participant in the previous iteration. A correct classification of a video resulted in a presentation of the given video with N frames less on the next iteration of the staircase, while a failure led to adding N frames more, where N decreased monotonously with the iteration number (n) following the formula N = Nframe/2 × 2-(n - 2)/2.5, where Nframe is the number of frames in the video. Iterations of the staircase were performed until N was smaller than 1 and hence the magnitude of up-down reversals became smaller than 1 frame, thus converging on that frame. The frame converged upon by the final iteration was taken as the frame on which the category (*Normal, How*, or *What*) becomes recognizable.

### Statistical Analyses

R 4.1.1 (R Core Team, 2020) was used for all statistical analyses. Participants’ ratings from the psychometric evaluation procedure were used for calculating validity (expressed as recognition accuracy in number and percentage of correctly classified videos) and inter-rater reliability (expressed as Fleiss’ *Kappa*; Fleiss, 1971). We performed an exploratory analysis investigating differences in recognition accuracy of videos from different categories (*Normal, How*, and *What*) and filming angles (portrait, left profile, and top) with 2-way within-subjects ANOVA.

Following the up-down staircase procedure, we investigated potential undesirable systematic effects of categories (*Normal, How*, and *What*) and angles (portrait, left profile, and top) on motion onset and timing of classification of videos, respectively with separate 2-way ANOVAs. This was deemed necessary as, although general differences in video properties across videos are expected, systematic differences between different groups of videos need to be brought to light so that future researchers can take them into account when designing experiments.

For all within-subjects ANOVAs, a Greenhouse-Geisser correction was applied when the Mauchly test indicated a significant departure from sphericity. Significant interactions were further analyzed by breaking down the main ANOVA into one-way ANOVAs, and simple effects were followed by post-hoc Tukey pairwise comparisons. The R script for performing all analyses can be found in the folder “All Data and Analyses” at https://osf.io/zexc4/.

## Results

### Validity and Reliability

Figure 4A presents participants’ overall category (*Normal, How*, or *What*) classification accuracy, which ranged between 85 and 135 videos (*Mean* = 119.63, *SD* = 10.41), corresponding to 63 % and 100 % accuracy respectively (*Mean* = 88.61 %, *SD* = 7.71 %). The average accuracy was significantly higher than that of 45 videos expected purely by chance (*t*_50_ = 51.18, *p* < 0.0001, 95% CI [116.69, 122.56], *d* = 7.17). Figure 4B presents the confusion matrix for classifications of *Normal, How*, and *What* videos.

**Figure 4.**
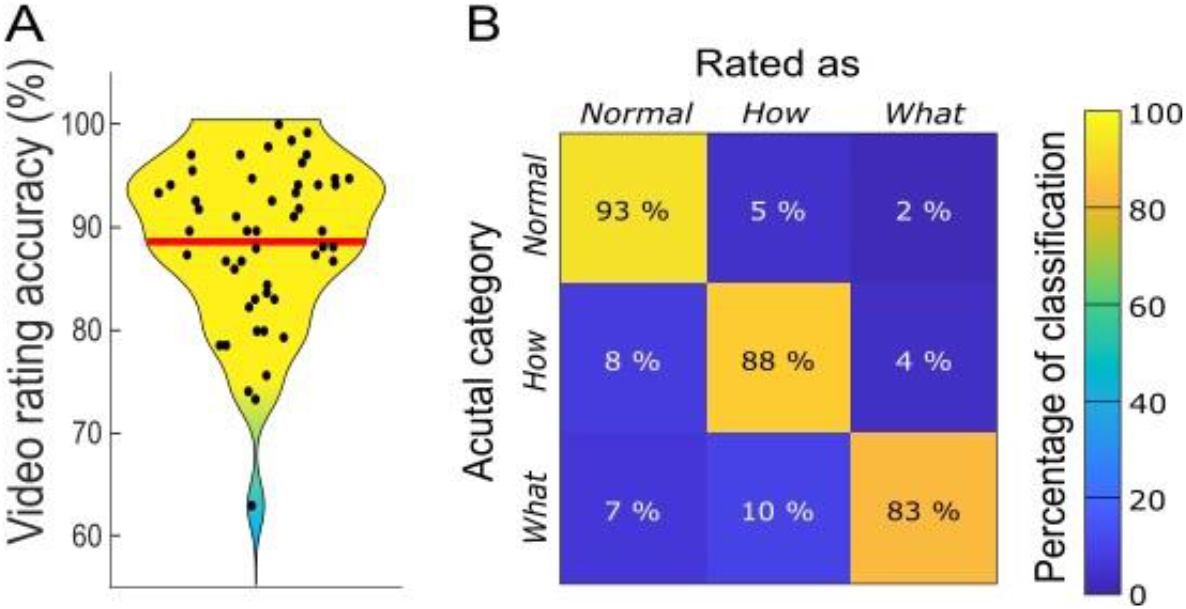
Video recognition accuracy of the whole sample (A) and recognition confusion matrix (B) Note. In (A) the red line indicates the mean recognition accuracy of 119.63 (88.61 %) videos.

It indicates that most misclassifications involved reporting *How* or *What* videos as *Normal*, and *What* videos as *How*. The proportion of misclassification was, however, quite small compared to the proportion of correct classifications, therefore establishing good validity of the present dataset.

The ANOVA investigating the extent to which the classification accuracy depended on videos’ category and filming angle revealed a significant effect of category (*F*_2, 98_ = 9.39, *p* < 0.001, partial η2 = 0.158), no effect of filming angle (*F*_2, 95_ = 0.55, *p* = 0.571, partial η2 = 0.011), and a significant category × filming angle interaction (*F*_4, 185_ = 3.08, *p* = 0.020, partial η2 = 0.058). To disentangle the interaction, we performed simple effects analyses of the factor category for each level of filming angle with one-way ANOVAs. These analyses revealed a significant effect of category on the classification accuracy for videos filmed from portrait angle (*F*_2, 98_ = 6.23, *p* = 0.003, η2 = 0.111), profile angle (*F*_2, 94_ = 12.11, *p* < 0.001, η2 = 0.195), and top angle (*F*_2, 100_ = 6.83, *p* = 0.002, η2 = 0.120). Figure 5 presents the outcome of post-hoc Tukey tests for each filming angle. A significant difference was found between *Normal* and *What* videos for all 3 angles, between *Normal* and *How* videos only within the top angle, and between *How* and *What* videos only within the profile angle. Considering all angles together, classification accuracy was marginally higher for *Normal* compared to *How* (*t*_50_ = 2.39, *p* = 0.052), significantly higher for *Normal* compared to *What* (*t*_50_ = 4.08, *p* = 0.0005), and marginally higher for *How* compared to *What* (*t*_50_ = 2.1, *p* = 0.099) videos. Although some statistically significant differences were found, it should be noted that values of classification accuracy were very high across all categories and angles, with mean accuracy ranging from 12.37 out of 15 for the *What* videos in profile view to 14.09 out of 15 for the *Normal* videos in profile view.

**Figure 5.**
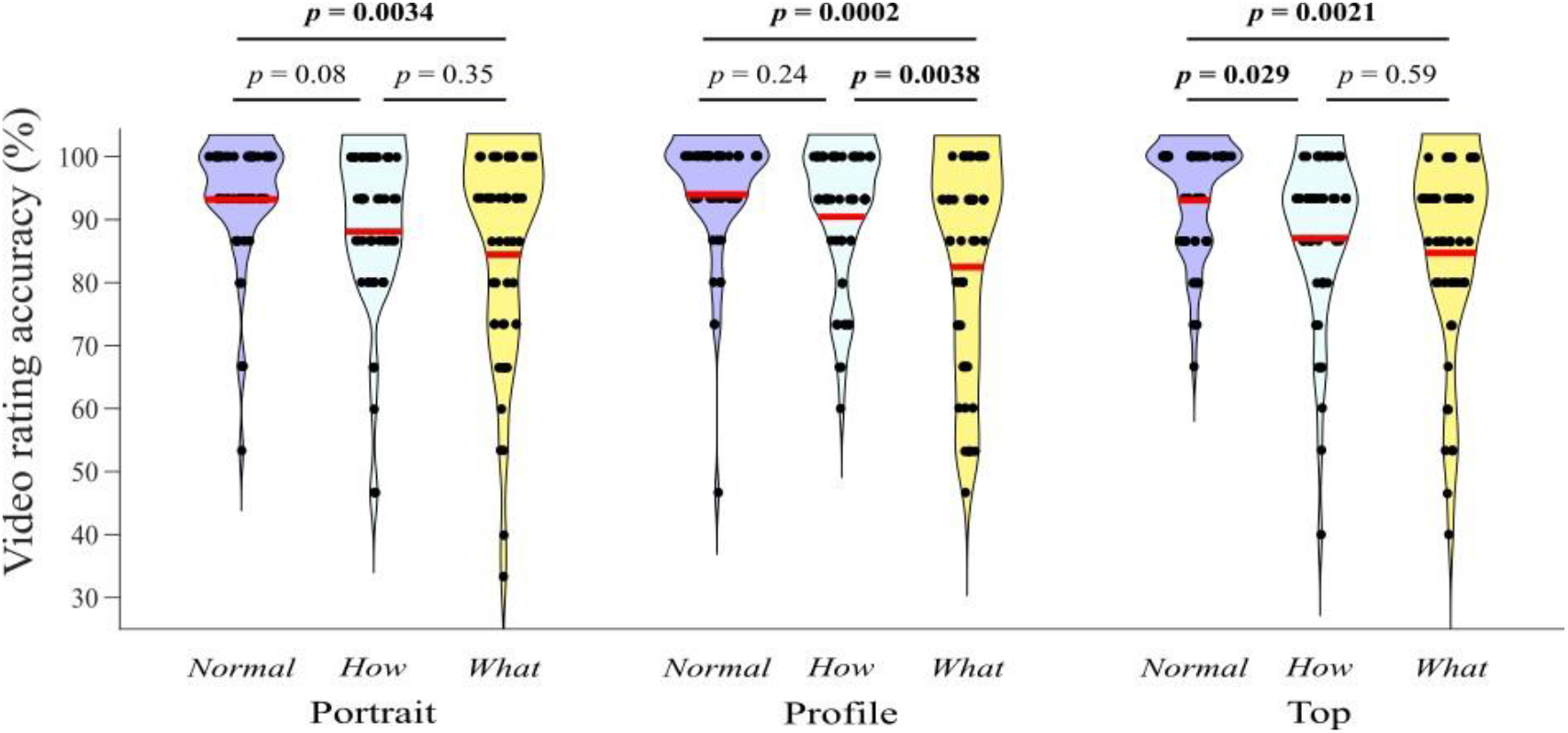
Distributions of the number of correctly recognized videos per category (Normal, How, and What) and filming angle (Portrait, Profile, and Top) Note. Red lines indicate the respective means. P-values correspond to Tukey tests.

### Motion Onset Detection

As expected, even with the strict detection parameters we specified, the motion detection system reported differences in the onset of motion across the videos, with motion onset detected in the range 2 – 18 frames (*Mean* = 2.5, *SD* = 1.79), corresponding to time 0.03 – 0.3 s (*Mean* = 0.04 s, *SD* = 0.03 s). The 2-way ANOVA investigating for differences in motion onset between videos from different category and filming angle revealed no effect of category (*F*_2, 126_ = 0.54, *p* = 0.583, partial η2 = 0.009), a significant effect of filming angle (*F*_2, 126_ = 4.4, *p* = 0.014, partial η2 = 0.065), and no category × filming angle interaction (*F*_4, 126_ = 1.06, *p* = 0.38, partial η2 = 0.032). Post-hoc Tukey tests revealed that in the portrait and top videos, motion was identified earlier than in the left profile videos (*t*_126_ = -2.65, *p* = 0.024 and *t*_126_ = -2.47, *p* = 0.039, respectively), with no difference in onset between portrait and top videos (*t*_126_ = -0.18, *p* = 0.982).

When comparing the output of the frame-differencing procedure to the motion onset frames identified by a human observer, we found a large degree of convergence with the exact same motion onset frame identified by the two for 113 (83.7 %) videos. A close inspection of the motion onset detection in the remaining 22 videos revealed that the discrepancy was as a result of instances of oversensitivity of the frame-differencing procedure which resulted in detection of small shadows, wrinkles in clothing, or eye movements of the actor. To circumvent this issue, as well as the discovered undesirable systematic differences in motion onset, we decided to edit all videos by cropping out the excessive still frames in the beginning and assuring that humanly perceptible motion onset begins exactly at frame 2 for every video. The success of this editing was verified by passing the videos to the frame-differencing algorithm and a visual inspection for the second time, which were in agreement for all 135 videos. The scripts for performing the edits of the videos are made available. The edited videos in .m4v and .mat format can be found in the folder “Video Database” at https://osf.io/zexc4/.

### Category Recognition

In just 17 between-subject iterations, the staircase procedure converged on a single frame within each video of the original database. The convergence frame was in the range 31 – 259 frames (*Mean* = 81, *SD* = 35.73), corresponding to time 0.52 – 4.32 s (*Mean* = 1.35 s, *SD* = 0.6 s). The 2-way ANOVA investigating for differences in the timing of category recognition between videos from different category and filming angle revealed a significant effect of category (*F*_2, 126_ = 5.01, *p* = 0.008, partial η2 = 0.074), no effect of filming angle (*F*_2, 126_ = 0.95, *p* = 0.39, partial η2 = 0.015) and no category × filming angle interaction (*F*_2, 126_ = 0.41, *p* = 0.799, partial η2 = 0.013). Figure 6B presents post-hoc Tukey tests on the factor category. These tests revealed that *Normal* videos were recognized on average 22.9 frames earlier than the *What* videos, with no significant difference between *Normal* and *How* and *How* and *What* videos.

**Figure 6.**
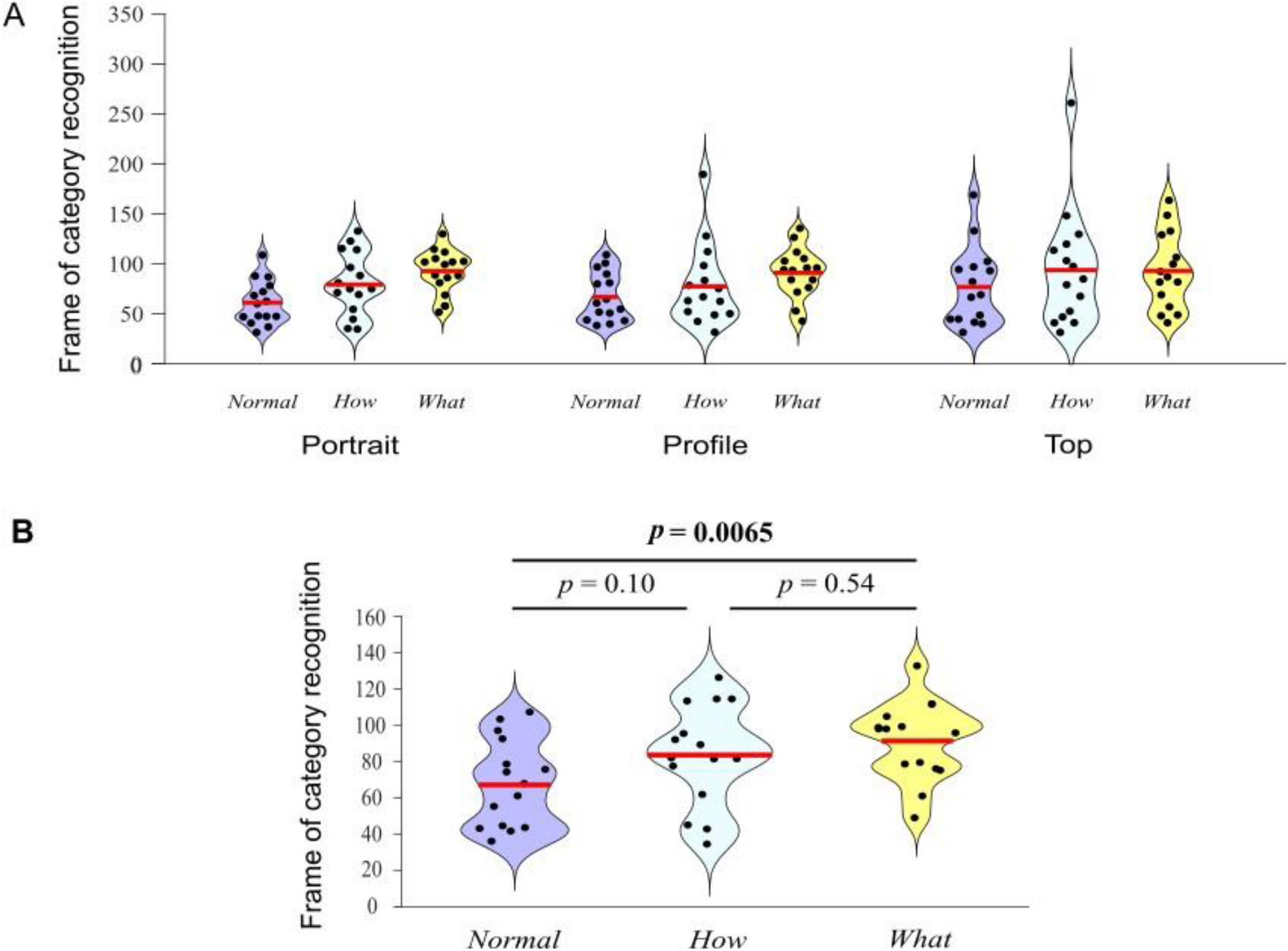
Distributions of the frames on which videos were recognized as Normal, How, or What for each category and filming angle (A) and Tukey tests for factor category (B).

This entire analysis investigating for differences in the timing of category recognition was repeated after the videos were edited to be equated for motion onset. The results of this analysis revealed a pattern of results identical to the one for the original video database. Each video’s frame of category recognition, before and after editing is presented in Appendix A.

## Discussion

The present article introduces a novel database of 135 video stimuli for action observation research in neuroimaging settings. It includes high definition videos of a male actor performing object-oriented actions from three categories: kinematically natural and goal-intact Normal actions, kinematically unnatural and goal-intact How actions, and kinematically natural and goal-violating What actions. This publicly available dataset will facilitate the investigation of both bottom-up and top-down neural processing of the characteristics of observed human actions.

Psychometric evaluation of the videos indicated high video classification accuracy and inter-rater agreement, conceptually translating to good validity and substantial reliability of the video categories. Although classification accuracy was very high for videos from all categories, we identified that classification accuracy was consistently higher for Normal compared to What videos in the portrait, profile, and top filming angles. While statistically significant, this difference consisted of, on average, one additional video in fifteen in the What category not being correctly classified, compared to the near perfect classification of Normal videos. Therefore, we refrain from interpreting it as a mental phenomenon of importance, or as a serious methodological challenge inherent in the use of our database. Instead, in line with the goal of the publication of our database, these findings could be the object of future psychophysics and neuroimaging investigations aimed at elucidating the recognition of different types of actions.

Subsequent to the psychometric evaluation, every video’s exact frame of motion onset was identified with a custom-made motion detection system. Some undesirable differences in motion onset of the original video database as well as slight inaccuracies in the frame-tracing procedure were discovered, therefore we re-edited all videos to assure that perceptible motion begins at exactly the second frame in each video. This assures that the Normal, How, and What videos in each filming angle are equivalent in terms of motion onset, which facilitates their presentation in neuroimaging experiments and circumvents the issue of jittered triggering altogether.

Moreover, the frame on which each video becomes discernibly recognizable as Normal, How, or What was also identified with a novel up-down staircase procedure. We identified significant differences in timing of category recognition between the Normal, How, and What videos with the Normal actions recognized 20–25 frames (or 0.33 s – 0.42 s) earlier than the How and What actions. Across videos, there were differences of up to ∼220 frames (or 3.7 s) in the timing of action category recognition. These results further highlight the importance of incorporating video motion characterization procedures into future experimental designs. Identifying the respective onset frames allows for counteracting the described differences by incorporating appropriately timed triggers. For instance, in addition to sending a trigger at the moment of presentation of each video on the screen, the experimenters will be able to send a second trigger corresponding to the exact moment of recognition of each video as identified by the staircase procedures. This approach would then allow for more precise grand averaging of epochs, precisely timed to the actual onset of the characteristic of interest. It is expected that such an approach will improve the sensitivity of event-related neuroimaging experiments. For this reason, the video database, the psychometric and motion evaluations, and the scripts required for performing these evaluations are made freely available for future scientific use. Moreover, our two motion evaluation procedures are well suited for characterizations of other already existing (Di Crosta et al., 2020; Umla-Runge et al., 2012; Urgen et al., 2022) or to-be-developed sets of video stimuli. Furthermore, any additional evaluations of the videos such as collection of arousal and valence ratings, normative validation in special populations, or collection of eye-tracking data are also greatly encouraged, as are any further edits of the videos. A fruitful direction for future research is a validation of the results of our motion onset frame-differencing algorithm and our category recognition up-down staircase with more advanced artificial intelligence and markerless motion capture methods to assure their precision of capturing nuanced human movements.

Our open-access video database greatly facilitates the process of stimulus preparation for subsequent action observation experiments. By relying on the present set of videos, researchers can design controlled investigations of the perception and comprehension of the kinematics of goal-directed human motion without investing resources into filming videos or depending on haphazard videos gleaned from the internet. Indeed, the reliance on standardized videos across experiments reduces the likelihood of introducing undesired variability into the experimental manipulation due to differences of stimuli. Therefore, reliance on the present set of videos could improve the comparability and replicability of action observation research within motor and cognitive neuroscience.

As it currently stands, the present dataset has a wide applicability for action observation research within various fields of inquiry. The videos were explicitly designed for application in electrophysiological (e.g., Electroencephalography and Magnetoencephalography), hemodynamic (functional Magnetic Resonance Imaging, Functional Near-Infrared Spectroscopy and Positron Emission Tomography), and transcranial stimulation (Transcranial Magnetic Stimulation, transcranial Direct Current Stimulation) studies of brain activity during action observation (e.g., Arnstein et al., 2011; Avanzini et al., 2012; Brunsdon, Bradford and Ferguson, 2019; Dinomais et al., 2013; Nedelko et al., 2010). However, they can also be utilized in investigations of the neural encoding of the kinematics of observed action (Bourguignon et al., 2013; Marty et al., 2018; Savaki et al., 2022), the goals and intentions of observed actions (Hamilton and Grafton, 2006; Nicholson, Roser and Bach, 2017), as well as the actions and goals afforded by objects (Bach, Bayliss and Tipper, 2011; Tipper, Paul and Hayes, 2006). Since our videos are highly recognizable, they can also be used in investigations of all above mentioned phenomena during action imitation and action learning interventions for clinical populations with impaired action comprehension such as autism spectrum disorder (Kaokhieo et al., 2023) and schizophrenia (Enticott et al., 2008). Such interventions are expected to strengthen the action perception–execution association, facilitate action embodiment (Tschacher, Giersch and Friston, 2017), and lead to improved social cognition. In addition, the videos from the Normal category can also serve as excellent training stimuli for action observation therapy – a promising intervention for motor recuperation in patients with movement disorders such as stroke (Ertelt et al., 2007; Fu et al., 2017) or Parkinson’s disease (Pelosin et al., 2010). The videos can also be used as naturalistic visual stimuli for cognitive neuroscience experiments investigating, among others, attention to and memorization of human action, supplementing the frequently used static pictures and non-naturalistic point-light displays (Gao, Bentin and Shen, 2015; Lu et al., 2016; Sifre et al., 2018). Further advances in all above-mentioned fields of research are necessary for gaining a more comprehensive understanding of the neural and cognitive processes which allow for perception and comprehension of observed action and, in turn, execution of appropriate corresponding motor reactions. Such understanding could foster training protocols for timely initial development and, in the case of acquired cognitive and motor impairments, effective recovery of these abilities – both crucial for the display of effective interpersonal behaviors within the cohesive structure of a healthy society.

## Conclusion

In closing, in hopes of facilitating action observation research in neuroimaging settings and encouraging open methodology, we provide a large open-access database of psychometrically evaluated videos of an actor performing movements explicitly designed to be kinematically correct or incorrect and goal-intact or goal-violating. We also provide the timing of action category recognition within each video for the sake of maximizing the precision of the assessment of the neural activity evoked by the observation of the different actions. The precise characterization of the videos in terms of psychometric properties and motion onset makes the present database a very suitable stimulus set for experiments aimed at investigating the neural correlates of perception, comprehension and acquisition of observed action.

## References

Arnstein, D. et al. (2011) ‘μ-suppression during action observation and execution correlates with BOLD in dorsal premotor, inferior parietal, and SI cortices’, The Journal of neuroscience: the official journal of the Society for Neuroscience, 31(40), pp. 14243–14249.

Avanzini, P. et al. (2012) ‘The dynamics of sensorimotor cortical oscillations during the observation of hand movements: an EEG study’, PloS one, 7(5), p. e37534.

Bach, P., Bayliss, A.P. and Tipper, S.P. (2011) ‘The predictive mirror: interactions of mirror and affordance processes during action observation’, Psychonomic bulletin & review, 18(1), pp. 171–176.

Beattie, G. and Shovelton, H. (2002) ‘An experimental investigation of some properties of individual iconic gestures that mediate their communicative power’, British Journal of Psychology, pp. 179–192.

Biagi, L. et al. (2016) ‘Action observation network in childhood: a comparative fMRI study with adults’, Developmental science, 19(6), pp. 1075–1086.

Bourguignon, M. et al. (2013) ‘Primary motor cortex and cerebellum are coupled with the kinematics of observed hand movements’, NeuroImage, 66, pp. 500–507.

Braadbaart, L., Williams, J.H.G. and Waiter, G.D. (2013) ‘Do mirror neuron areas mediate mu rhythm suppression during imitation and action observation?’, International journal of psychophysiology: official journal of the International Organization of Psychophysiology, 89(1), pp. 99–105.

Bradski, G.R. and Kaehler, A. (2008) Learning OpenCV: Computer Vision with the OpenCV Library.

Brunsdon, V.E.A., Bradford, E.E.F. and Ferguson, H.J. (2019) ‘Sensorimotor mu rhythm during action observation changes across the lifespan independently from social cognitive processes’, Developmental cognitive neuroscience, 38, p. 100659.

Buccino, G. et al. (2004) ‘Neural circuits underlying imitation learning of hand actions: an event-related fMRI study’, Neuron, 42(2), pp. 323–334.

Caillaud, M. et al. (2020) ‘Influence of emotional complexity on the neural substrates of affective theory of mind’, Human brain mapping, 41(1), pp. 139–149.

Cheng, C.-H. et al. (2017) ‘Differential motor cortex excitability during observation of normal and abnormal goal-directed movement patterns’, Neuroscience research, 123, pp. 36– 42.

Di Crosta, A. et al. (2020) ‘The Chieti Affective Action Videos database, a resource for the study of emotions in psychology’, Scientific data, 7(1), p. 32.

Dinomais, M. et al. (2013) ‘Effect of observation of simple hand movement on brain activations in patients with unilateral cerebral palsy: an fMRI study’, Research in developmental disabilities, 34(6), pp. 1928–1937.

Enticott, P.G. et al. (2008) ‘Reduced motor facilitation during action observation in schizophrenia: a mirror neuron deficit?’, Schizophrenia research, 102(1-3), pp. 116–121.

Ertelt, D. et al. (2007) ‘Action observation has a positive impact on rehabilitation of motor deficits after stroke’, NeuroImage, 36 Suppl 2, pp. T164–73.

Fleiss, J. L. (1971). ‘Measuring nominal scale agreement among many raters’, Psychological Bulletin, 76(5), pp. 378–382.

Fu, J. et al. (2017) ‘Effects of action observation therapy on upper extremity function, daily activities and motion evoked potential in cerebral infarction patients’, Medicine, 96(42), p. e8080.

Gao, Z., Bentin, S. and Shen, M. (2015) ‘Rehearsing biological motion in working memory: an EEG study’, Journal of cognitive neuroscience, 27(1), pp. 198–209.

Gera, G. et al. (2010) ‘Motor abundance contributes to resolving multiple kinematic task constraints’, Motor control, 14(1), pp. 83–115.

Hamilton, A.F. de C. and Grafton, S.T. (2006) ‘Goal representation in human anterior intraparietal sulcus’, The Journal of neuroscience: the official journal of the Society for Neuroscience, 26(4), pp. 1133–1137.

Huettel, S.A. (2012) ‘Event-related fMRI in cognition’, NeuroImage, 62(2), pp. 1152–1156.

Iacoboni, M. et al. (2005) ‘Grasping the intentions of others with one’s own mirror neuron system’, PLoS biology, 3(3), p. e79.

Kaernbach, C. (1991) ‘Simple adaptive testing with the weighted up-down method’, Perception & psychophysics, 49(3), pp. 227–229.

Kaokhieo, J. et al. (2023) ‘Effects of repetitive transcranial magnetic stimulation combined with action-observation-execution on social interaction and communication in autism spectrum disorder: Feasibility study’, Brain research, 1804, p. 148258.

Levitt, H. (1971) ‘Transformed up-down methods in psychoacoustics’, The Journal of the Acoustical Society of America, 49(2), p. Suppl 2:467+.

Lu, X. et al. (2016) ‘Holding Biological Motion in Working Memory: An fMRI Study’, Frontiers in human neuroscience, 10, p. 251.

Malfait, N. et al. (2010) ‘fMRI activation during observation of others’ reach errors’, Journal of cognitive neuroscience, 22(7), pp. 1493–1503.

Marty, B. et al. (2018) ‘Movement Kinematics Dynamically Modulates the Rolandic ∼ 20-Hz Rhythm During Goal-Directed Executed and Observed Hand Actions’, Brain topography, 31(4), pp. 566–576.

Moriguchi, Y. et al. (2009) ‘The human mirror neuron system in a population with deficient self-awareness: an fMRI study in alexithymia’, Human brain mapping, 30(7), pp. 2063–2076.

Muthukumaraswamy, S.D. and Johnson, B.W. (2004) ‘Primary motor cortex activation during action observation revealed by wavelet analysis of the EEG’, Clinical neurophysiology: official journal of the International Federation of Clinical Neurophysiology, 115(8), pp. 1760–1766.

Nedelko, V. et al. (2010) ‘Age-independent activation in areas of the mirror neuron system during action observation and action imagery. A fMRI study’, Restorative neurology and neuroscience, 28(6), pp. 737–747.

Nicholson, T., Roser, M. and Bach, P. (2017) ‘Understanding the Goals of Everyday Instrumental Actions Is Primarily Linked to Object, Not Motor-Kinematic, Information: Evidence from fMRI’, PloS one, 12(1), p. e0169700.

Orban, G.A., Ferri, S. and Platonov, A. (2019) ‘The role of putative human anterior intraparietal sulcus area in observed manipulative action discrimination’, Brain and behavior, 9(3), p. e01226.

Paxton, A. and Dale, R. (2013) ‘Frame-differencing methods for measuring bodily synchrony in conversation’, Behavior research methods, 45(2), pp. 329–343.

Pelosin, E. et al. (2010) ‘Action observation improves freezing of gait in patients with Parkinson’s disease’, Neurorehabilitation and neural repair, 24(8), pp. 746–752.

Platonov, A. and Orban, G.A. (2016) ‘Action observation: the less-explored part of higher-order vision’, Scientific reports, 6, p. 36742

Pütten, A.M.R. der et al. (2014) ‘Investigations on empathy towards humans and robots using fMRI’, Computers in Human Behavior, pp. 201–212.

Quandt, L.C. and Marshall, P.J. (2014) ‘The effect of action experience on sensorimotor EEG rhythms during action observation’, Neuropsychologia, 56, pp. 401–408.

Ramseyer, F.T. (2020) ‘Motion energy analysis (MEA): A primer on the assessment of motion from video’, Journal of counseling psychology, 67(4), pp. 536–549.

Ramseyer, F. and Tschacher, W. (2011) ‘Nonverbal synchrony in psychotherapy: coordinated body movement reflects relationship quality and outcome’, Journal of consulting and clinical psychology, 79(3), pp. 284–295.

Savaki, H.E. et al. (2022) ‘Action Observation Responses Are Influenced by Movement Kinematics and Target Identity’, Cerebral cortex, 32(3), pp. 490–503.

Saxe, R. et al. (2004) ‘A region of right posterior superior temporal sulcus responds to observed intentional actions’, Neuropsychologia, 42(11), pp. 1435–1446.

Scott, M.W. et al. (2020) ‘Motor imagery during action observation enhances imitation of everyday rhythmical actions in children with and without developmental coordination disorder’, Human movement science, 71, p. 102620.

Sifre, R. et al. (2018) ‘A Longitudinal Investigation of Preferential Attention to Biological Motion in 2-to 24-Month-Old Infants’, Scientific reports, 8(1), p. 2527.

Sonkusare, S., Breakspear, M. and Guo, C. (2019) ‘Naturalistic Stimuli in Neuroscience: Critically Acclaimed’, Trends in cognitive sciences, 23(8), pp. 699–714.

Spengler, S., Bird, G. and Brass, M. (2010) ‘Hyperimitation of actions is related to reduced understanding of others’ minds in autism spectrum conditions’, Biological psychiatry, 68(12), pp. 1148–1155.

Stapel, J.C. et al. (2010) ‘Motor activation during observation of unusual versus ordinary actions in infancy’, Social neuroscience, 5(5-6), pp. 451–460.

Steen, M.C. van der, van der Steen, M.C. and Bongers, R.M. (2011) ‘Joint angle variability and co-variation in a reaching with a rod task’, Experimental Brain Research, pp. 411–422.

Suzuki, S. and Abe, K. (1985). ‘Topological structural analysis of digitized binary images by border following’, Computer Vision, Graphics, and Image Processing, 30(1), pp. 32–46.

Sylwester, K. et al. (2012). ‘The role of Theory of Mind in assessing cooperative intentions’, Personality and Individual Differences, 52(10), pp. 113–117

Tholen, M.G. et al. (2020) ‘Functional magnetic resonance imaging (fMRI) item analysis of empathy and theory of mind’, Human brain mapping, 41(10), pp. 2611–2628.

Tipper, S.P., Paul, M.A. and Hayes, A.E. (2006) ‘Vision-for-action: the effects of object property discrimination and action state on affordance compatibility effects’, Psychonomic bulletin & review, 13(3), pp. 493–498.

Trettenbrein, P.C. and Zaccarella, E. (2021) ‘Controlling Video Stimuli in Sign Language and Gesture Research: The Package for Analyzing Motion-Tracking Data in’, Frontiers in psychology, 12, p. 628728.

Tschacher, W., Giersch, A. and Friston, K. (2017) ‘Embodiment and Schizophrenia: A Review of Implications and Applications’, Schizophrenia bulletin, 43(4), pp. 745–753.

Umla-Runge, K. et al. (2012) ‘An action video clip database rated for familiarity in China and Germany’, Behavior research methods, 44(4), pp. 946–953.

Urgen, B.A. et al. (2022) ‘A Large Video Set of Natural Human Actions for Visual and Cognitive Neuroscience Studies and Its Validation with fMRI’, Brain sciences, 13(1), pp. 61.

Vrigkas, M., Nikou, C. and Kakadiaris, I. A. (2015). ‘A review of human activity recognition methods. Frontiers in Robotics and AI, 2.

Wetherill, G.B. and Levitt, H. (1965) ‘Sequential estimation of points on a psychometric function’, The British journal of mathematical and statistical psychology, 18, pp. 1–10.

Zouba, N. et al. (2008). ‘Monitoring activities of daily living (ADLs) of elderly based on 3D key human postures’, ICVW 2008 Cognitive Vision, pp. 37–50.

